# Structural model of Cyc2, the primary electron acceptor of *Acidithiobacillus ferrooxidans*’ respiratory chain, as a modular cytochrome - β-barrel fusion protein, and mechanistic proposals based on this model

**DOI:** 10.1101/262287

**Authors:** Luciano A. Abriata

## Abstract

*Acidithiobacillus ferrooxidans* oxidizes Fe(II) to Fe(III) to feed electrons into its respiratory chain. The primary electron acceptor of this complex system is Cyc2, an outer membrane protein of unknown structure. This work proposes a feasible model of Cyc2’s global structure, based on homology modeling, residue-residue coevolution data, bioinformatics predictions and limited knowledge about Cyc2’s function. The proposal is that the sequence segment spanning residues ~30 to ~90 folds as a cytochrome-like domain that contains a heme group which would presumably bind and oxidize external Fe(II), whereas the remaining segment from residue ~90 until the end adopts a β-barrel fold similar to that of most outer membrane proteins. Such model differs strongly from a published model, but is backed up by more data and is more compatible with the known topology of outer membrane proteins and with Cyc2’s function of internalizing reducing equivalents. The small size of the cytochrome-like domain would allow it to reside inside, and/or slide through, the β-barrel domain, thus communicating in a controlled fashion the extracellular medium with the periplasm to import electrons through the outer membrane. All the models discussed are provided as PyMOL session files in the Supporting Information and can be visualized online at http://lucianoabriata.altervista.org/modelshome.html

---

Respiration in *Acidithiobacillus ferrooxidans* proceeds through oxidation of Fe(II) to Fe(III) and oxygen reduction in a complex electron transfer chain.^1,2^ The primary electron acceptor of this system is Cyc2, an outer membrane protein that most likely delivers electrons to rusticyanin.^3,4^

As many membrane proteins, Cyc2 is difficult to handle in the laboratory and so far no structures have been solved for it or for homologs that could be used as templates for homology modeling. This work reports structural predictions for *A. ferrooxidans* Cyc2 (as declared in Uniprot entry B7JAQ7_ACIF2) based on homology modeling guided by residue-residue coevolution analysis and sequence-based analyses. The emerging model suggests a modular architecture consisting of a signal peptide, a small heme-binding cytochrome-like domain, a functionally relevant flexible linker and a large transmembrane β-barrel domain. Under such model the cyt-like domain could reside inside, or even slide back and forth through, the β-barrel domain to obtain Fe(II) from the extracellular medium and oxidize it to deliver its electron to downstream electron transport proteins like rusticyanin, in the periplasmic space.

## Results and Discussion

### Sequence-based predictions of secondary structure, disorder and domain structure

As annotated in Uniprot, Cyc2 is predicted to have an N-terminal signal peptide spanning residues 1 to 31, followed by a domain almost 460 residues long that does not match any known protein family at least as defined in PFAM. Furthermore, BLAST searches to the Protein Data Bank do not retrieve any proteins of similar sequence with known structures.

Secondary structure prediction by PSIPRED (Fig. 1A) propose an N-terminal α-helix that matches the purported signal peptide, followed by three additional, shorter helices and then by multiple stretches of β-strands. Between two of the three small helices, there is a CAACH motif like that of heme-binding motifs in cytochromes. At this point a modular architecture seems to appear that consists of a signal peptide followed by a small cytochrome-like domain and then a β-sheet-rich domain (Fig. 1B). According to disorder predictions, the small cytochrome-like and the β-sheet-rich domains are separated by a very flexible linker around 20 residue long (centered at around residue 120, Fig. 1C). Two other smaller stretches of flexible residues might be relevant at around residue 275 and 425.

**Figure 1.**
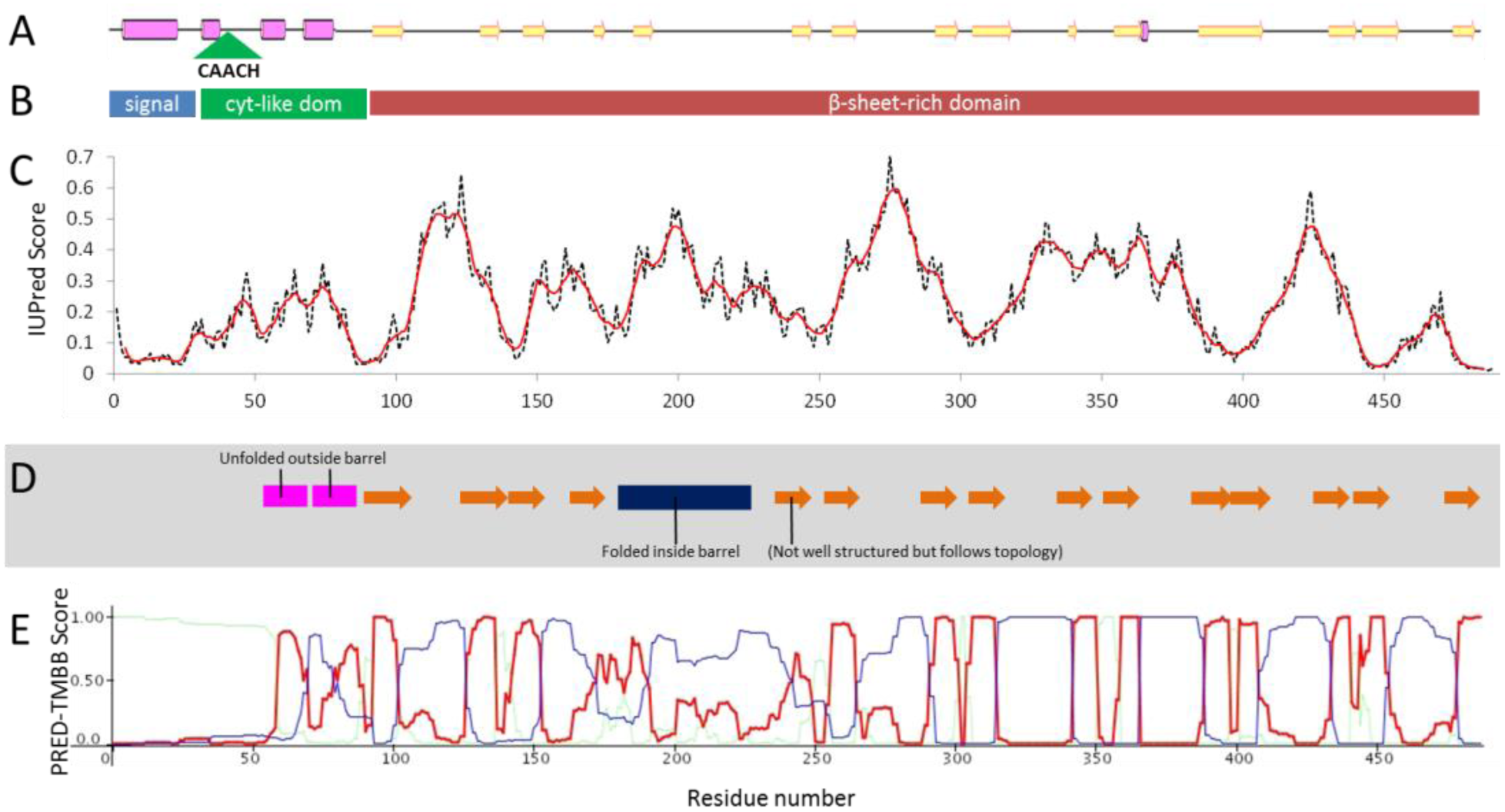
(A) PSIPRED secondary structure prediction (pink boxes are ct-helices, yellow arrows are P-strands) and the heme-binding CAACH motif. (B) Proposed domain topology. (C) Disorder prediction from IUPred (dashed is raw data, red line is 7 residue average). (D) Secondary structures observed in the proposed structural model (orange arrows are P-strands of the P-barrel). (E) Prediction of P-barrel topology by PREDTMBB (high scores in red trace) and localization (high scores in blue trace predict external localization, low values internal).

### Structural modeling of Cyc2 from weak homology to templates, assisted by contacts from pairwise residue coevolution analysis

BLAST searches of Cyc2’s sequence to the Protein Data Bank return weak matches, including partial matches to outer membrane β-barrel proteins and also to other proteins of radically different fold. Therefore, modeling of Cyc2 is anticipated to be hard.

Running Cyc2’s sequence on the I-TASSER^5^ server returns models of very different topologies (Figure 2A). Model #1 displays the topology of the Adhesive Tip pilin GBS104 from Group B *Streptococcus agalactiae*, visually somewhat similar to the structural model proposed for Cyc2 in ^6^. Such a model would imply that Cyc2 does not span the full membrane width, which is expected given its function. On the other hand, I-TASSER’s models #2, #3 and #5 display features of β-barrel folds, as expected for outer membrane proteins. Model #4 shows yet another fold, different from that of the models #1, 2, 3 and 5.

**Figure 2.**
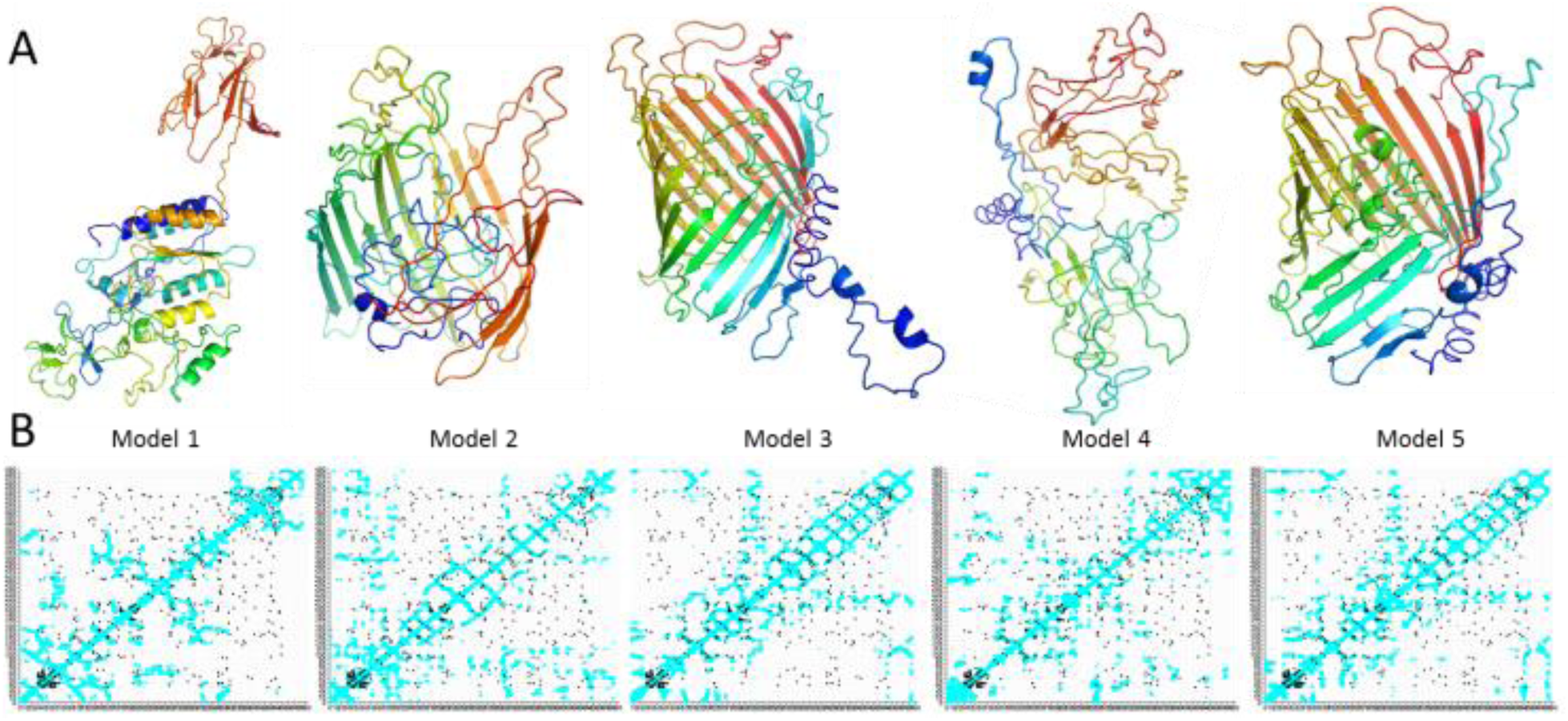
(A) The five models returned by I-TASSER for the full Cyc2 sequence (colored from blue at the N-terminus to red at the C-terminus). (B) Contact maps of models (cyan) compared to the pairs of residues that display coevolution coupling > 0.02 (black, Gremlin values). In the plots in (B) both x and y axes are Residue numbers.

By comparing residue-residue coevolution strengths computed from an alignment of Cyc2-like proteins with the Gremlin server,^7,8^ it appears that the predicted contacts match best with the contacts present in model #3 (Fig. 2B and different cutoffs of the coevolution data in Fig. 3). An additional I-TASSER run augmented with Gremlin’s pairs of coevolving residues at p>0.025 (Fig. 3, top) returns only models with β-barrel folds, all similar to each other. Of these, the top model (#1) is considered here as the final model. This model (Fig. 4A,B) improves slightly from model #3 of the first I-TASSER run and matches the coevolution data marginally better. Considering that the match is not perfect, and the many distortions observed in the model, it is expected that it only accounts for overall shape and topology but not for details like the exact ends of the beta strands and loop conformations.

**Figure 3.**
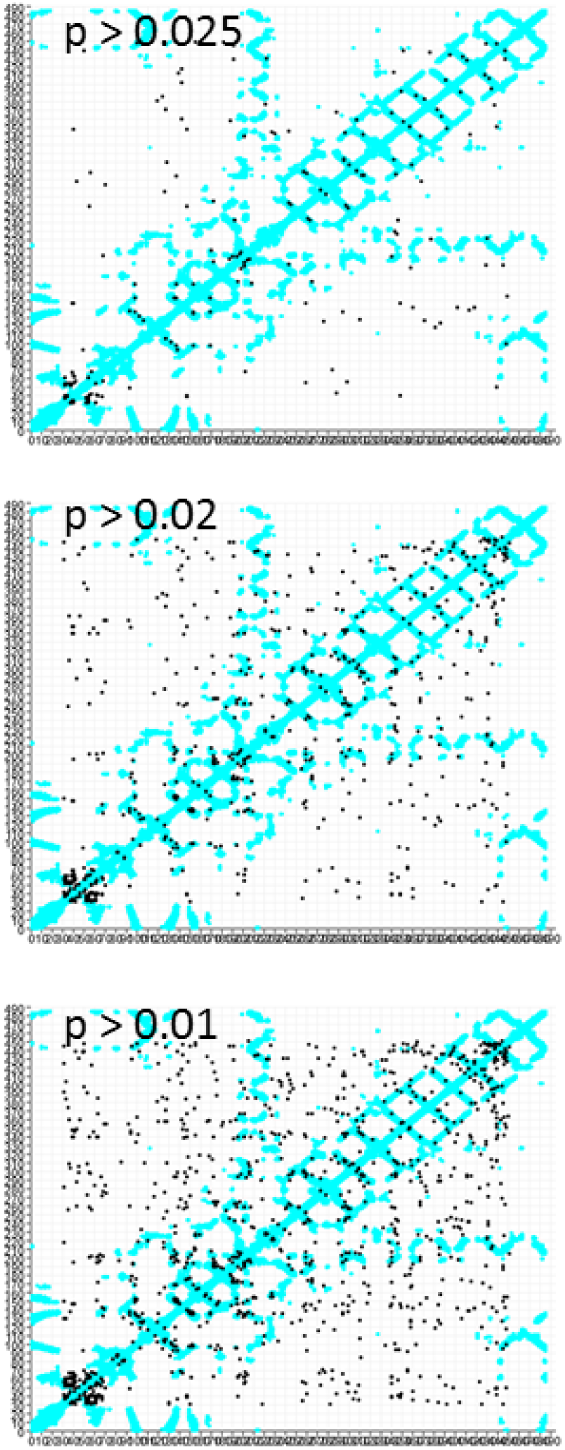
Comparison of Model 3’s contact map (cyan) with the pairs of coevolving residues at different cutoffs of coevolution strength (black dots).

**Figure 4.**
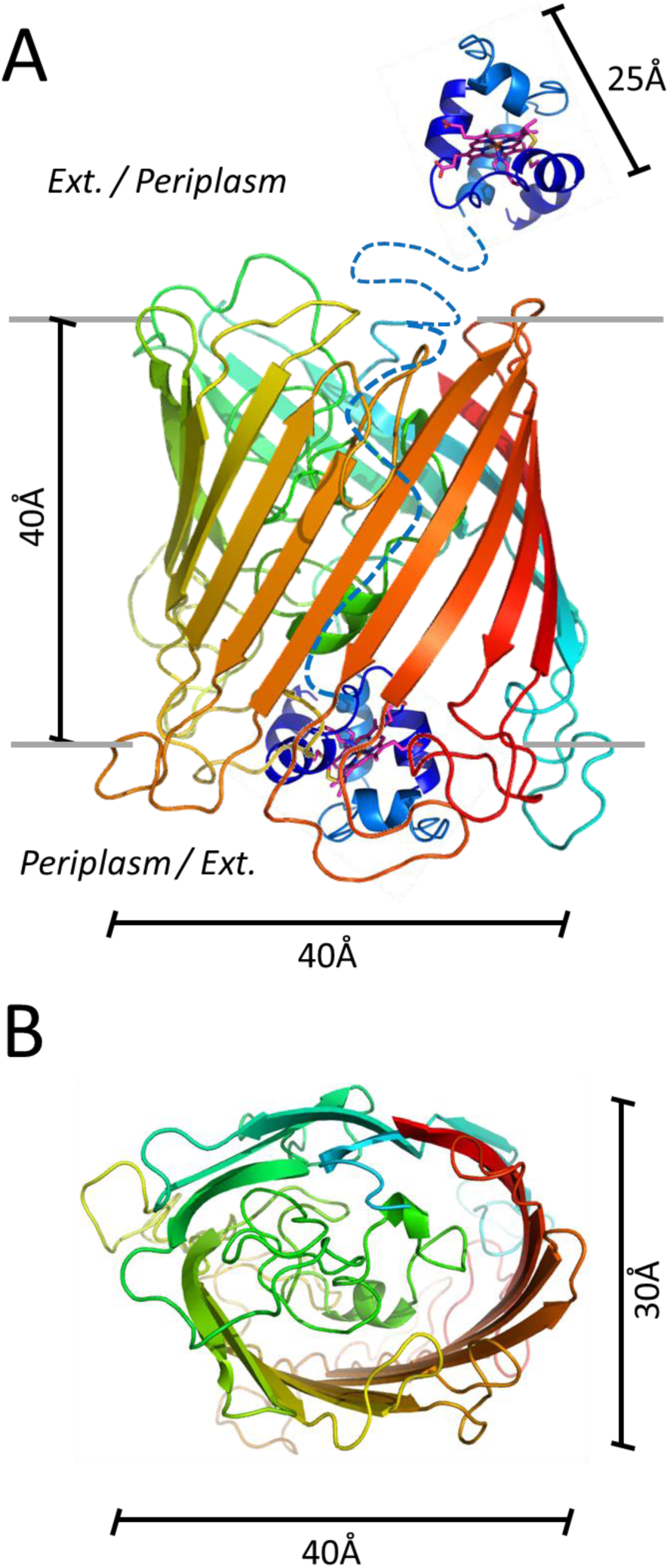
(A) Side views of models of Cyc2’s cytochrome-like and β-barrel domains, connected by a flexible linker shown as dashes. The backbone trace is colored from blue at the N-terminus to red at the C-terminus. The cytochrome-like domain is shown in two possible localizations: exposed to the periplasm or outer cell space, or docked inside the β-barrel domain. (B) Top view of the model of the β-barrel domain. Approximate sizes are shown in both panels.

The β-strands of the final model shown in Fig. 4 (mapped onto the amino acid sequence in Fig. 1D) correspond quite well to those predicted by PSIPRED from Cyc2’s sequence (Fig. 1A). Furthermore, having observed that the protein likely adopts a β-barrel fold prompts prediction of this topology from sequence with a specialized server. Such prediction by the PREDTMBB server (Fig. 1E) also matches reasonably, although not perfectly, with the topology of β-strands in the β-barrel domain of the model.

### Structural features of the Cyc2 structural model and functional relevance of these features

Overall, this simple strategy puts forward a very draft model of Cyc2’s topology (Fig. 4), which, despite surely having multiple deficiencies, can help guide future biochemical and structural studies. Essentially, according to the model, Cyc2 consists of a β-barrel as most other known outer membrane proteins, with an N-terminal extension that likely contains a small cytochrome-like with a heme group (Fig. 4A).

A side view of the final model of the β-barrel domain shows dominance of hydrophobic amino acids in the external side of the barrel, as expected for an integral membrane protein. Along the membrane normal the protein is around 40 Å long, indicating it would span the full membrane. On the membrane plane the model looks like an oval of major and minor diameters of around 40 and 30 Å, respectively.

The small cytochrome-like domain modelled with I-TASSER from homology to PDB ID 2zzs (Fig. 4A) shows most hydrophobic residues in its core and polar residues at its surface. Its three main sizes are around 25, 20 and 20 Å, therefore it could fit inside the polar interior of the β-barrel domain, either statically or with the freedom to slide through the barrel exchanging between localizations in the extracellular region, inside the β-barrel, and/or in the periplasm. In the former, “static” case Fe(II) would enter the barrel and deliver the electrons at its base on the periplasmic side, where downstream electron-accepting proteins should dock. In the latter, “dynamic” scenario, the cytochrome-like domain could swim around the barrel to fish Fe(II) ions from the exterior and then slide through the barrel toward the periplasm to import the reduced ion.

It is important that none of the two purported electron acceptors from Cyc2, *i.e.* rusticyanin and Cyc4, would fit inside Cyc2’s barrel domain. Rusticyanin’s major sizes are around 40, 37 and 33 Å, while Cyc4’s are around 45, 30 and 25 Å. They should therefore dock on the periplasmic side of Cyc2, or interact with its cytochrome-like domain if it can slide completely through the β-barrel to reach the periplasm.

## Conclusions

Combining different sources of structural information led here to a new model of Cyc2’s structure, radically different to that previously proposed.^6^ The modular domain organization proposed here suggests a heme-containing cytochrome-like domain linked through a flexible linker to an outer membrane-embedded β-barrel. The cytochrome-like domain would be posed to capture Fe(II) or its electrons from the extracellular medium, and the β-barrel domain would mediate communication of the cytochrome-like domain with the downstream electron carriers located in the periplasm.

## Supporting Information

A short methods section, as well as PyMOL session files for the “final” model of Cyc2’s β-barrel domain (produced with I-TASSER guided by Gremlin coevolution constraints) and for the model of the cytochrome-like domain (aligned in space with its template to highlight the location of the heme group) are provided in the Supporting Information. Models can also be inspected in interactive 3D online at http://lucianoabriata.altervista.org/modelshome.html

## Acknowledgement

I acknowledge Dr. Marianne Ilbert (Aix-Marseille University) for comments and discussion.

